# The emerging contaminant Cl-PFESA/F-53B is toxic to meiotic cell division and reproduction in *Arabidopsis thaliana*

**DOI:** 10.1101/2025.11.03.686200

**Authors:** Yuting Chen, Xueying Cui, Ziming Ren, Huiqi Fu, Yufeng Luo, Linji Xu, Ziwei Song, Yonghua Qin, Guanghui Yu, Xiaoning Lei, Bing Liu

## Abstract

The emerging contaminant Cl-PFESA/F-53B damages vegetative development in plants and has been evidenced to be toxic to both mitotic and meiotic cells in animals. We have recently reported that F-53B induces nuclei unviability in meristematic cells leading to disrupted root development in Arabidopsis (*Arabidopsis thaliana*). However, the toxicity of F-53B to reproductive system in plants remains unelucidated. In this study, by using cytogenetic and microscopic approaches, we analyzed embryo and anther development and multiple meiosis processes in Arabidopsis exposed to 50 or 100 μM F-53B. We showed that F-53B disrupts embryo development which results in reduced seed setting in Arabidopsis. Histochemical staining of anthers and a live-imaging assay using a reporter line that expresses GFP-tagged Aborted Microspores (AMS), a key regulator of the tapetum, demonstrated that F-53B impairs anther development. Moreover, we showed that F-53B interferes with chromosome segregation and/or distribution and alters microtubule organization during male meiosis. Quantification of the chiasmata and immunolocalization of the Human Enhancer of Invasion 10 (HEI10) protein on diakinesis chromosomes suggested that F-53B reduces crossover rate. Furthermore, F-53B reduced the number of DNA Meiotic Recombinase 1 (DMC1) protein foci on zygotene chromosomes in Arabidopsis wild-type Columbia-0 (Col-0), and it partially promoted meiotic chromosome integrity in Arabidopsis depleted with *Ataxia Telangiectasia Mutated* (*ATM*), a central regulator of DNA double-strand break (DSB) repair, which suggested that F-53B reduces meiotic DSB formation. Taken together, our study reveals the toxicity of F-53B to meiotic cells and reproduction in Arabidopsis, which highlights its potential threats to agricultural safety and ecological diversity.

**Highlight:**

- F-53B impairs embryo and anther development in Arabidopsis.
- F-53B interferes with chromosome segregation and distribution during male meiosis.
- F-53B interferes with meiotic microtubule organization.
- F-53B reduces meiotic crossover formation.
- F-53B reduces DNA double-strand breaks in meiocytes.

## 1 Introduction

Reproductive development is crucial for the completion of the life cycle and fertility in flowering plants. At early stage of microsporogenesis, pollen mother cells (PMCs) undergo meiosis which involves twice nuclei division after one round of chromosomal DNA replication to generate microspores (Zickler and Kleckner, 2023). At prophase I, homologous chromosomes (homologs) pair and synapse and undergo recombination to exchange DNA materials through crossing-overs (COs). Meiotic recombination is initiated by the programmed formation of double-strand breaks (DSBs), which are catalyzed by a conserved type II topoisomerase (topoisomerase VI, subunit A) SPO11 together with other proteins in the DSB machinery (Grelon et al., 2001; Stacey et al., 2006; Vrielynck et al., 2021). DSBs are mainly repaired via the homologous recombination pathway, in which the recombinases RAD51 and DMC1 play an indispensable role (Li et al., 2004; Sanchez-Moran et al., 2007). RAD51 acts downstream of the conserved kinase Ataxia Telangiectasia Mutated (ATM) to repair DSBs using sister chromatids as the templates; DMC1 accessorized by RAD51 drives DSB repair using homologs as the templates (Chen et al., 2021; Cloud et al., 2012; Da Ines et al., 2013; Kurzbauer et al., 2012; Lan et al., 2020; Pradillo et al., 2012; Sanchez-Moran et al., 2007; Yao et al., 2020). In plants, dysfunction of DMC1 leads to impaired CO formation and univalents, while lesions in RAD51 function cause severe chromosome fragmentation (Da Ines et al., 2013). Despite a large number of DSBs is generated on meiotic chromosomes, only a small fraction of them is repaired into COs, which are classified into type I and II based on the pathways of the involved recombinases (Wang and Copenhaver, 2018; Zelkowski et al., 2019). Type I COs, which occupies the most proportion of COs, are catalyzed by the proteins (e.g., HEI10 and MLH1) acting in the ZMM (ZIPs, MER3, and MSH4/5) pathway. Type I COs are spaced along the chromosomes under the regulation of mechanism called CO interference (Ziolkowski, 2023). Type II COs are mediated by structure-selective endonucleases such as MUS81 (Hartung et al., 2006; Hollingsworth and Brill, 2004), and are not subject to CO interference. For each pair of homologs, at least one CO must be formed to ensure faithful segregation of homologs (Jones and Franklin, 2006). Despite the significances for genome stability and genetic diversity, meiosis in plants is sensitive to the alterations in environmental conditions, for example, temperature and light (Fu et al., 2024; Lei et al., 2020; Liu et al., 2019; Ning et al., 2021). Studies in many plant species have revealed that meiosis is one of the most stress-sensitive periods during reproductive development (Bomblies et al., 2015; De Storme and Geelen, 2014; Liu et al., 2019). In addition, meiosis is non-cell-autonomously regulated by somatic tissues especially the tapetum in the anthers (Lei and Liu, 2020; Tidy et al., 2022).

Per- and polyfluoroalkyl substances (PFASs) are anthropogenic chemicals manufactured in the 1950s and have been widely used in various industrial and products due to chemical and thermal stability (Gaines, 2023; Glüge et al., 2020). However, the release of PFASs during production and daily use has led to pollutions in the main environmental matrices (Brusseau et al., 2020; De Silva et al., 2021; Nakayama et al., 2019). Perfluorooctane sulfonic acid (PFOS) is the most common PFAS and has been detected in various environmental compartments, raising serious concerns for global crisis of PFOS contamination (Wee and Aris, 2023b, a). Increasing studies have reported that PFOS is absorbable by plants, which triggers stress responses and defects in plant growth and development (Adu et al., 2023; Chen et al., 2020; Chi et al., 2024; Ghisi et al., 2019; He et al., 2023; Li et al., 2022; Pietrini et al., 2024; Song et al., 2024; Wang et al., 2020). Chlorinated polyfluoroalkyl ether sulfonate (Cl-PFESA/F-53B, F-53B is used later on for simplification), which primarily comprises 6:2 Cl-PFESA and 8:2 Cl-PFESA, has been specifically used in China as the main PFOS alternative in the electroplating industry for decades (Wang et al., 2013).

F-53B has been detected globally raising serious concerns for an ecological contamination (Ti et al., 2018). F-53B has been found to inhibit development in several plant species by triggering DNA damages and/or cell unviability (Li et al., 2023; Lin et al., 2020; Pan et al., 2021; Qiu et al., 2024; Qu et al., 2010). Compared with PFOS, F-53B reveals a higher accumulation ability in soil and plant tissues (Li et al., 2020; Lin et al., 2020; Xu et al., 2022; Zhang et al., 2021) and a higher toxicity likely by triggering higher levels of hydroxyl free radicals (Li et al., 2023; Lin et al., 2020; Pan et al., 2021). However, our recent study uncovered that F-53B does not induce reactive oxygen species (ROS) but instead damages cell viability and division in meristematic cells in Arabidopsis seedlings via a different pathway as ROS does (Zhao et al., 2025). Moreover, in both mammals and several plant species, F-53B has been found or proposed to trigger ectopic DSB formation (Li et al., 2023; Rogakou et al., 1998; Wang et al., 2015). While, we have provided evidence by multiple approaches in a previous study that F-53B does not induce DSB formation in Arabidopsis root cells (Zhao et al., 2025). These findings indicate both conserved and divergent effects of F-53B on the development in organisms, which reveals a complexity of its biotoxicity.

A recent study reported that F-53B interferes with chromosome segregation and spindle organization during female meiosis in mouse (Chu et al., 2025), revealing the toxicity of F-53B to sexual reproductive system in mammals. However, the potential impacts of F-53B on reproductive development in plants remain unelucidated. In the present study, we assessed the effects of F-53B on multiple reproductive development processes in Arabidopsis (*Arabidopsis thaliana*). We report that F-53B interferes with meiotic cell division and attenuates CO formation. Moreover, we show that F-53B disrupts tapetum and embryo development in Arabidopsis. Our study sheds lights on the toxicity of F-53B to the sexual-reproducing system in plants, which highlights its potential threats to agricultural safety and ecological diversity.

## 2 Material and Methods

### 2.1 Plant materials and growth conditions

*Arabidopsis thaliana* (L.) accession Columbia-0 (Col-0) was used as the wild-type. The *atm-2* mutant (SALK_006953) (Zhao et al., 2023) and the *pAMS::AMS-GFP* reporter (Xiong et al., 2016) were used in this study. Genotyping of *atm-2* was performed using the primers: ‘atm-2 F’ (ATCCATGTGGTTCAGTCTTGC) and ‘atm-2 R’ (TTGGTATCCTGCAGAGGAAAG). Seeds were germinated and cultivated in half-strength Murashige & Skoog (1/2 MS) medium in a growth chamber with a 16 h day/8 h night, 20°C, and 50% humidity condition following vernalization in dark and 4°C conditions for 3 days.

### 2.2 Treatment of F-53B

Young flowering Arabidopsis individuals were cut at terminal position of their main shoots and were immersed in a 50 or 100 μM F-53B solution that have been shown to induce nuclei damage leading to vegetative development abortion in Arabidopsis (Zhao et al., 2025). This treatment enables transmission of chemicals in the solution into flower tissues via the transpiration flow in the inflorescence (Armstrong, 2013). The samples used for chromosome spreading and immunolocalization assays were isolated at 48 h post the exposure to F-53B. The F-53B product, which comprises 97% 6:2 Cl-PFESA and 3% 8:2 Cl-PFESA (Wu et al., 2023), was purchased from Jianglai Biotechnology Co., Ltd (Shanghai, China).

### 2.3 Analysis of silique and embryo development

To determine the impact of F-53B on silique development, the developed siliques on the main shoots were removed before the plants being immersed in control or F-53B solutions. At the 7^th^ day post exposure to F-53B, newly-developed siliques were isolated and counted, in which developing embryos were analyzed.

### 2.4 Alexander staining of the anthers

Alexander staining was performed as described previously (Alexander, 1969; Fu et al., 2024). Newly-developed flower buds, which were about to flower, from the 1^st^ to 7^th^ day post exposure to control or F-53B solutions were isolated for Alexander staining.

### 2.5 Preparation of meiotic chromosome spreads

Preparation of meiotic chromosome spreads was performed by referring to (Ross et al., 1996). Inflorescences of young Arabidopsis were fixed in precooled Carnoy’s fixative for at least 24 h. Meiosis-staged flower buds were washed twice with distilled water and once with citrate buffer (10 mM, pH = 4.5), followed by incubation in a digestion enzyme mixture (0.3% pectolyase and 0.3% cellulase in citrate buffer, 10 mM, pH = 4.5) at 37°C for 2.5 h. Subsequently, the digested flower buds were washed once in distilled water and macerated in distilled water on a glass slide. Two aliquots of 60% acetic acid were added to the slide, which was dried on a hotplate at 45°C for 2 min. The slide was washed with ice-cold Carnoy’s fixative and then air-dried. DAPI (5 μg/ml) diluted in antifade mounting medium was added to the slide, and the coverslip was mounted and sealed with nail polish. The number of chromosome fragments in metaphase I, anaphase I, metaphase II, and telophase II meiocytes was calculated as previously described (Zhao et al., 2023).

### 2.6 Immunolocalization

Immunolocalization of meiotic proteins and tubulin was performed by referring to (Zhao et al., 2024). The antibodies against HEI10 (rabbit) (Fu et al., 2022), DMC1 (rabbit) (Ning et al., 2021) and α-tubulin (rat) (Lei et al., 2020) were diluted by 1:200; the antibodies against SYN1 (rabbit) (Fu et al., 2022) and SYN1 (mouse) (Zhao et al., 2023) were diluted by 1:500 and 1:150, respectively. The secondary antibodies have been described previously (Zhao et al., 2024).

### 2.7 Live-imaging of the reporter

To analyze the expression of AMS, the anthers at different developmental stages in flowering Col-0 expressing the *pAMS::AMS-GFP* reporter (Xiong et al., 2016) at the 1^st^ to 6^th^ day post exposure to F-53B were isolated and placed on a glass slide with a drop of distilled water being added to the samples, which was mounted with a cover slide and examined under an inverted fluorescence microscope.

### 2.8 Quantification of fluorescent foci and fluorescence intensity

The numbers of HEI10 and DMC1 protein foci on meiotic chromosomes were quantified as previously described (Zhao et al., 2023). Only the fluorescent foci merged onto DAPI-stained chromosome bodies were counted.

### 2.9 Microscopy

Fluorescence images (excitation wavelength peaks from 496 to 553 nm; emission wavelength peaks from 519 to 568 nm) were recorded using an Olympus IX83 inverted fluorescence microscope with a X-Cite lamp and a Prime BSI camera. Bifluorescent images and Z-stacks were processed by Image J.

### 2.10 Statistics

The significance levels were determined based on unpaired *t*-tests or Chi-squared tests with GraphPad Prism (v.8). The numbers of analyzed cells and/or biological replicates are shown in figures or legends; variation bars indicate standard deviation (SD).

## 3 Results

### 3.1 Reduced fertility in Arabidopsis exposed to F-53B due to aborted embryo development

To assess the toxicity of F-53B to reproduction in plants, we evaluated the impact of F-53B on silique development in Arabidopsis (*Arabidopsis thaliana*). Young flowering Arabidopsis accession Columbia-0 (Col-0) plants were exposed to F-53B via a method that has been used for 5-Ethynyl-2’-deoxyuridine (EdU) or Bromodeoxyuridine (BrdU) labelling of DNA in meiocytes (Armstrong et al., 2003; Hernández Sánchez-Rebato et al., 2024; Sanchez-Moran et al., 2007). Col-0 with cut terminal main shoots were immersed in a 50 or 100 μM F-53B solution, which have been reported to induce nuclei damage leading to vegetative development abortion in Arabidopsis (Zhao et al., 2025). This treatment enables transmission of chemicals in the solution into flower tissues via the transpiration flow in the inflorescence (Armstrong, 2013). The number of newly-developed siliques in Col-0 exposed to F-53B or a control solution at the 7^th^ day post treatment was compared, which revealed that Col-0 exposed to either 50 or 100 μM F-53B had significantly less newly-developed siliques than that in control (Fig. 1A-D; CK vs. 50 μM F-53B, *P* < 0.01; CK vs. 100 μM F-53B, *P* < 0.001). In addition, the siliques yielded by plants exposed to F-53B were shorter than those in control (Fig. 1A-C, yellow arrows; E), which suggested a reduced fertility. We checked embryo development in the siliques and found that most embryos (95.5%) in control siliques have developed and showed similar sizes (Fig. 1F and I). However, Col-0 exposed to 50 or 100 μM F-53B showed that only 66.7% and 51.5% embryos, respectively, have developed, yet smaller than those in control (Fig. 1G-I). The aborted embryos showed a small and/or withered configuration (Fig. 1G and H, red arrows). These findings suggested that F-53B damages embryo development in Arabidopsis.

**Figure 1.**
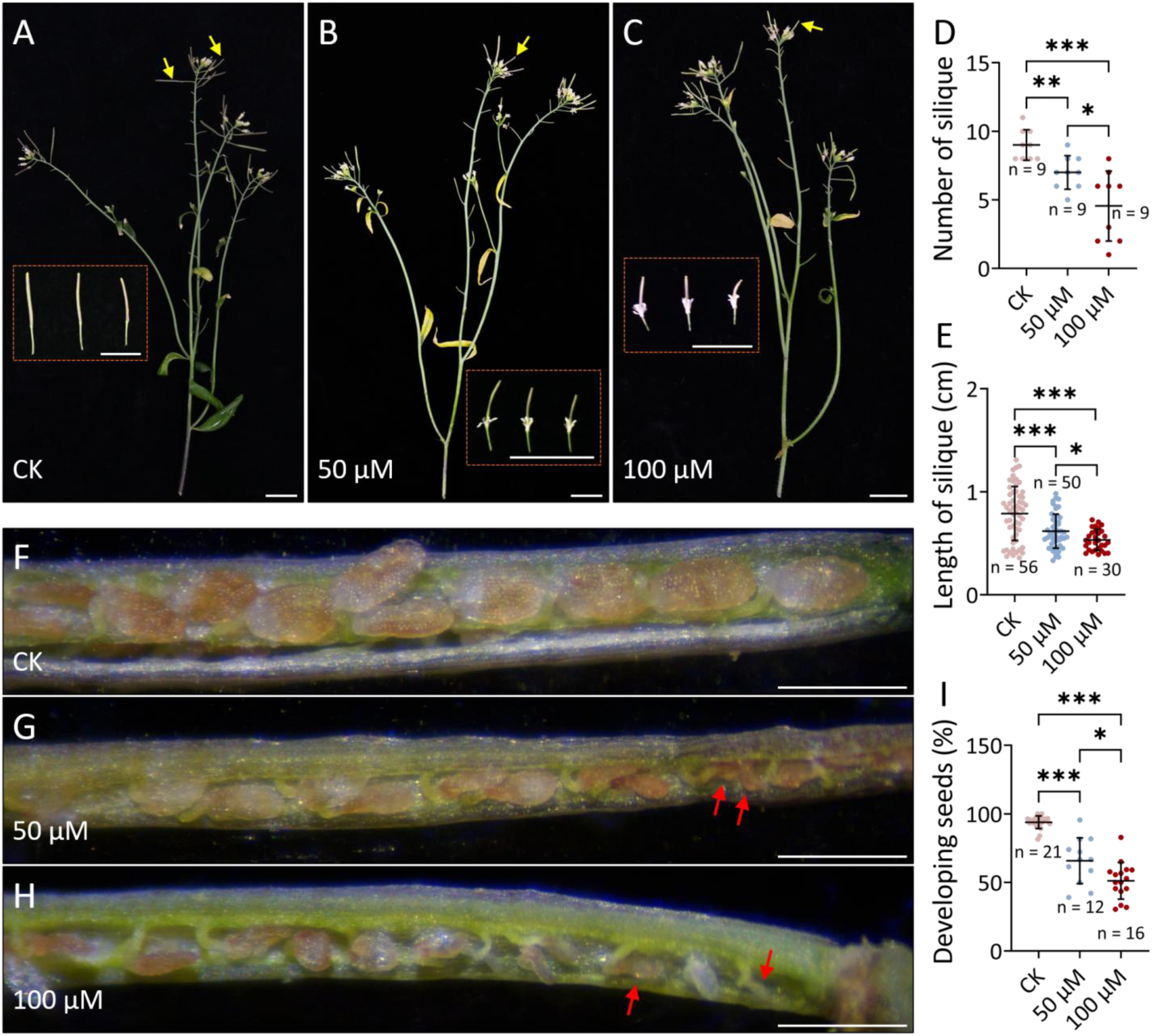
F-53B impairs reproduction in Arabidopsis. A-C, Col-0 plants under control conditions (A) and after exposure to 50 (B) or 100 μM (C) F-53B for 7 days. The yellow arrows indicate newly-developed siliques during the treatment; a close-up view for each condition is shown in the brown boxes. Scale bars = 1 cm. D and E, Graphs showing the number (D) and length (E) of the newly-developed siliques from Col-0 plants under control conditions or after exposure to F-53B for 7 days. F-H, Representative images of developing and aborted embryos in siliques produced by Col-0 plants under control conditions (F) and after exposure to 50 (G) or 100 μM (H) F-53B for 7 days. The red arrows indicate aborted embryos. Scale bars = 500 μm. I, Graph showing the frequency of developing embryos in newly-developed siliques from Col-0 plants under control conditions or after exposure to F-53B for 7 days. The significance levels were determined based on unpaired *t* tests; n indicates the number of plant individuals (D) or siliques (E and I); *** indicates P < 0.001; ** indicates P < 0.01; * indicates P < 0.05.

### 3.2 F-53B disrupts gametogenesis and tapetum development

To explore the impacts of F-53B on gametogenesis in Arabidopsis, we performed Alexander staining of the anthers following 1-to 6-day post the exposure to F-53B. At 1-to 3-day post F-53B treatment, there was not obvious difference in viable pollen grain formation between control and F-53B-exposed plants (Fig. 2A). On the 4^th^ day, however, anthers with aborted pollen grains were observed in plants exposed to 100 μM F-53B (Fig. 2B). On the 5^th^ and 6^th^ day, plants exposed to either 50 or 100 μM F-53B showed small and shriveled anthers, in which almost no viable pollen grain was found (Fig. 2C and D). Based on these findings we conclude that F-53B impairs male gametogenesis.

**Figure 2.**
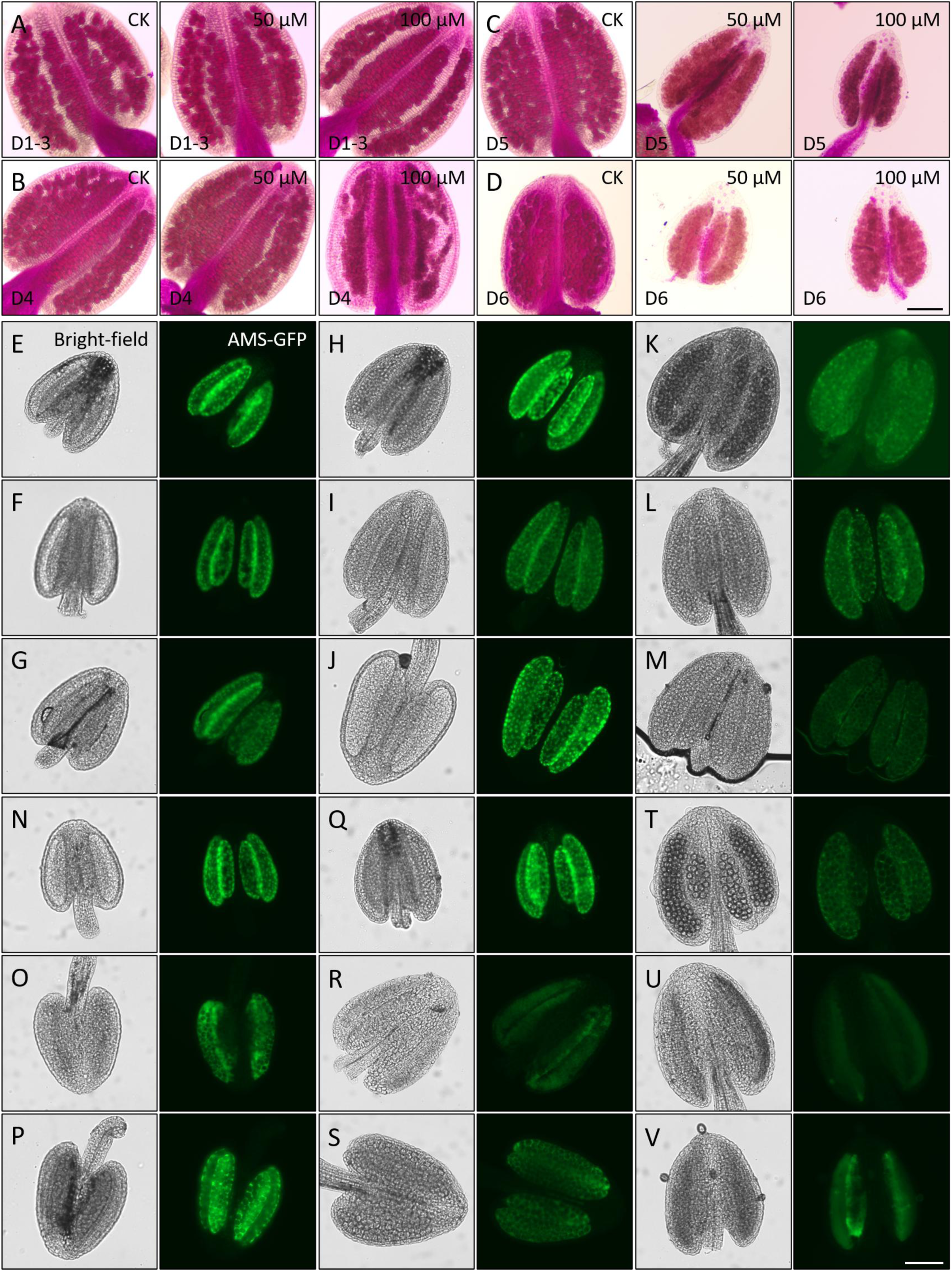
F-53B impairs gametogenesis. A-D, Alexander staining of the anthers in control and at 1 to 6-day post exposure to 50 and 100 μM F-53B. D1-D6, day 1-6. Scale bar = 100 μm. E-V, Live-imaging of the *pAMS::AMS-GFP* reporter expressing in anthers at tetrad (E-G, N-P), uni-cellular (H-J, Q-S) and bi- and/or tri-cellular (K-M, T-V) microspore stages in control (E, H, K, N, Q and T) or after the exposure to 50 (F, I, L, O, R and U) and 100 μM (G, J, M, P, S and V) F-53B for 48 (E-M) or 72 h (N-V). Scale bar = 50 μm.

Since tapetum plays a crucial role in assuring faithful male gametogenesis (Scott et al., 2004), we monitored the impact of F-53B on tapetum by analyzing the expression of *Aborted Microspores* (*AMS*), which regulates tapetum development and programmed cell death by acting in the conserved Dysfunctional Tapetum 1 (DYT1)-Tapetal Development and Function 1 (TDF1) cascade (Feng et al., 2012; Ferguson et al., 2017; Gu et al., 2014; Xu et al., 2010). We performed live-imaging using a GFP-tagged AMS reporter, *pAMS::AMS-GFP* (Xiong et al., 2016), which shows sensitivity to UV stress (Fu et al., 2024), at the 2^nd^ and 3^rd^ day post exposure to F-53B. Under normal conditions, AMS was specifically expressed in the tapetum with the anthers at the tetrad, uni-cellular and bi-or tri-cellular microspore stages (Fig. 2E, H and K). In comparison, the anthers at bi-or tri-cellular microspore stage showed reduced AMS-GFP signals because of the programmed cell death (PCD) in the tapetal cells (Fig. 2E, H and K), which is essential for microspore development (Alonso-Peral et al., 2010; Ferguson et al., 2017; Gu et al., 2014; Li et al., 2011; Xu et al., 2010). At 48 h post exposure to either 50 or 100 μM F-53B, the reporter did not show an obvious difference in the expression pattern of AMS-GFP (Fig. 2F and G, I and J, L and M). At 72 h post treatment, however, both the plants exposed to 50 or 100 μM F-53B exhibited reduced AMS-GFP signals at tetrad stage; at uni-cellular and bi- and/or tri-cellular microspore stages, almost no AMS-GFP signal could be detected in the anthers (Fig. 2N-V), which suggested that tapetum PCD ectopically occurred, or the development and/or function of tapetum was impaired.

### 3.3 Arabidopsis exposed to F-53B shows defects in meiotic chromosome segregation and distribution

To explore whether early processes during microsporogenesis, i.e., meiosis, is subject to the exposure to F-53B, we monitored behaviors of chromosomes during male meiosis in Col-0 by analyzing chromosome spreads stained by 4’,6-diamidino-2-phenylindole (DAPI). In control flowers, meiocytes at pachytene stage showed fully paired and synapsed homologous chromosomes (homologs), which developed into five bivalents at diakinesis and metaphase I (Fig. 3A-C). At anaphase I, homologs separated with each other to the opposite cell poles (Fig. 3D) and were aligned at metaphase II cell plates ready for the segregation of sister chromatids (Fig. 3E). Four chromosome sets were isolated at telophase II (Fig. 3F), which developed into four haploid nuclei at the tetrad stage, indicating the completion of male meiosis (Fig. 3G). In Col-0 exposed to 50 or 100 μM F-53B, homolog pairing and synapsis at pachytene occurred regularly (Fig. 3H and P), and most analyzed diakinesis and metaphase I meiocytes showed five bivalents (Fig. 3I and Q; J and S). However, we occasionally observed univalents at diakinesis and/or metaphase I stages (Fig. 3K, R, T and U, red arrows) under both conditions, which suggested that the assurance of CO formation was interfered. No defects in homolog separation were detected at anaphase I (Fig. 3L and V), possibly due to the low percentage of the defects in CO formation. At metaphase II, however, aberrant chromosome alignment and distribution occurred at 22.8% and 17.2%, respectively, in Col exposed to 50 or 100 μM (Fig. 3M, W and A’) F-53B. At the end of meiosis II, approximately 4.9% and 11.1% meiocytes in 50 or 100 μM F-53B-treated Col-0, respectively, displayed irregular chromosome and nuclei segregation and/or distribution (Fig. 3N and X, normal; O, Y and Z, abnormal), which will likely lead to aneuploid gametes. Overall, these data suggest that F-53B interferes with chromosome segregation and distribution in Arabidopsis meiosis.

**Figure 3.**
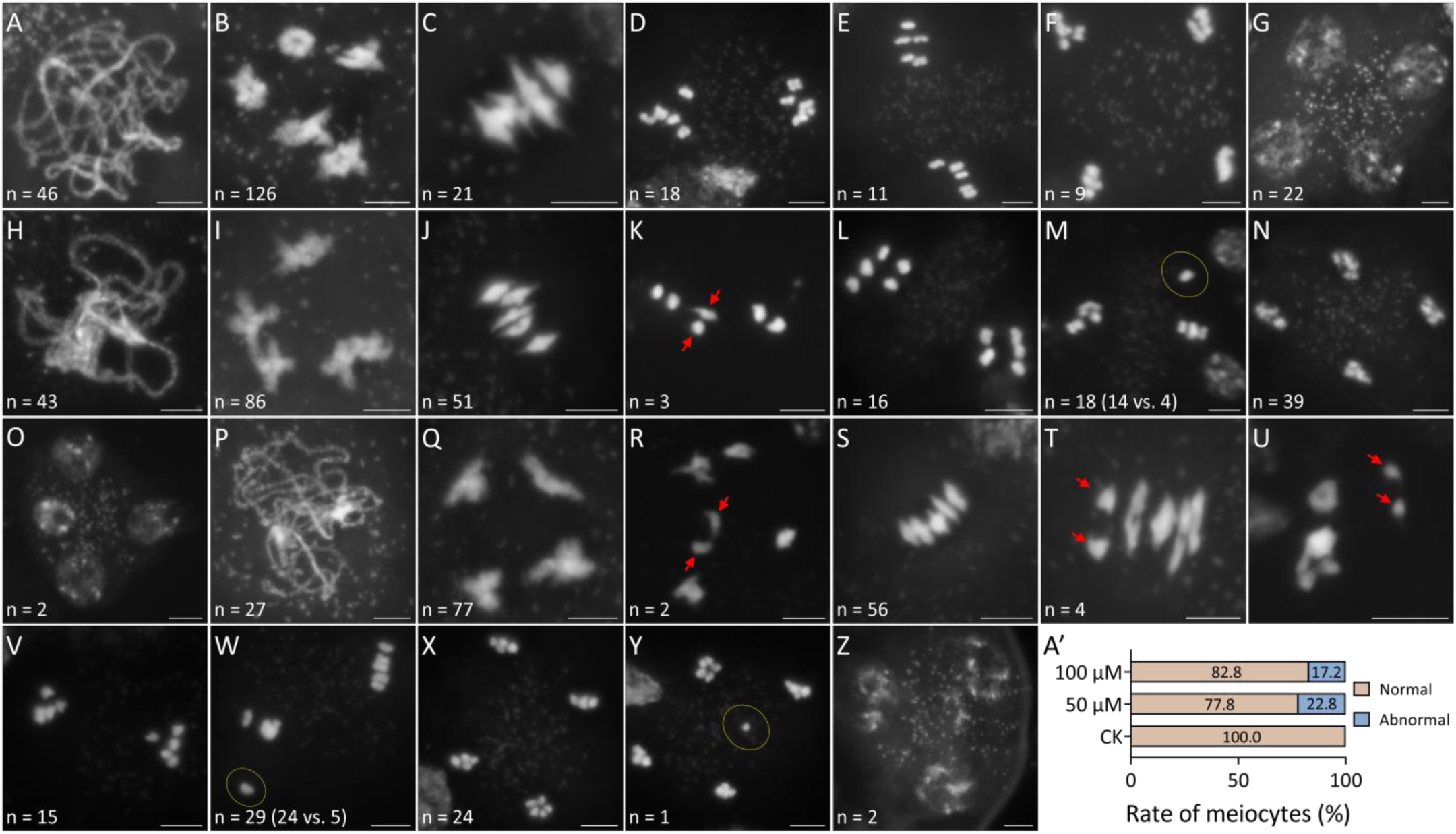
Chromosome behaviors during male meiosis in Col-0 exposed to F-53B. A-Z, DAPI-staining of meiocytes at pachytene (A, H and P), diakinesis (B, I, Q and R), metaphase I (C, J, K, S-U), anaphase I (D, L and V), metaphase II (E, M and W), telophase II (F, N, X and Y) and tetrad (G, O and Z) stages in Col-0 under control conditions (A-G), or exposed to 50 (H-O) or 100 μM (P-Z) F-53B for 48 h. The red arrows indicate univalents; yellow circles indicate irregularly distributed chromosomes. In the panels M and W, the numbers of meiocytes showing normal chromosome behaviors vs. the numbers of meiocytes showing abnormal chromosome behaviors were indicated in the parentheses. n indicates the number of analyzed meiocytes; scale bars = 5 μm. A’, Graph showing the fraction of meiocytes showing normal or abnormal chromosome distribution at telophase II in Col-0 under control conditions, or exposed to 50 or 100 μM F-53B for 48 h.

### 3.4 F-53B interferes with spindle and phragmoplast organization during meiosis

In mouse, F-53B interferes with the organization of microtubules during female meiosis (Chu et al., 2025). Our recent study revealed that F-53B damages microtubule-dependent cell plate formation in meristematic cells at root tips in Arabidopsis (Zhao et al., 2025). To determine whether microtubules in meiotic cells in plants is also subject to F-53B toxicity, we performed immunolocalization of α-tubulin and analyzed microtubule organization in Col-0 under control conditions or exposed to 50 or 100 μM F-53B. In control plants, chromosomes during prophase I were wrapped by a microtubular network, which was organized into a spindle structure at metaphase I with microtubules attaching to the kinetochores of the chromosomes that were aligned at the central region of the cell (Supplemental Fig. S1A; Fig. 4A). At anaphase I and interkinesis, homologs were separated to the opposite cell poles, between which a phragmoplast structure was built (Supplemental Fig. S1B and C). At metaphase II, two spindles were constructed to drive the segregation of sister chromatids, and at telophase II, two phragmoplasts were built between the newly-separated sister chromosome sets (Fig. 4B and C). At the end of male meiosis, radial microtubule arrays were assembled between four haploid nuclei to facilitate cytokinesis, which leaded to production of a tetrad (Supplemental Fig. S1D).

**Figure 4.**
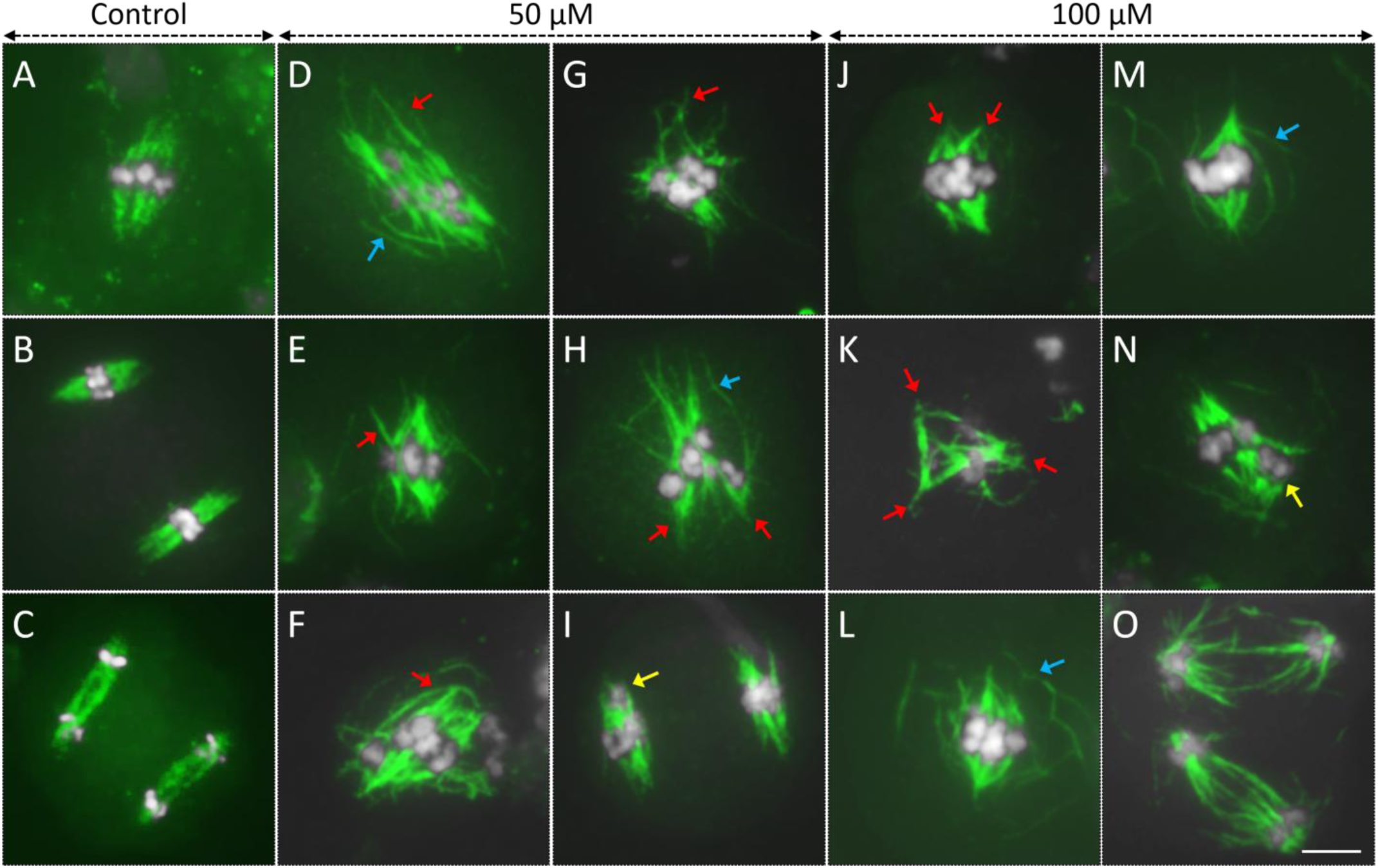
Arabidopsis exposed to F-53B shows alterations in microtubule organization during meiosis. A-O, Immunolocalization of α-tubulin in meiocytes at metaphase I (A, D-H, J-N), metaphase II (B and I), and telophase II (C and O) stages in Arabidopsis under control conditions (A-C), or exposed to 50 (D-I) or 100 μM (J-O) F-53B. White, DAPI; green, α-tubulin. The red arrows indicate abnormal α-tubulin organization or morphology; blue arrows indicate irregularly-assembled microtubule filaments; yellow arrows indicate defectively-distributed chromosomes. The scale bar applies to all panels in this figure and indicates 5 μm.

In meiocytes of Col-0 exposed to 50 or 100 μM F-53B, no obvious alteration in microtubule organization was visualized during prophase I (Supplemental Fig. S1E and N). However, at metaphase I, defects in spindle morphology and/or organization were observed in Col-0 exposed to either F-53B concentration (Supplemental Fig. S1F and O, normal; Fig. 4D-H and J-N, abnormal), which included: 1), mis-orientation of microtubules that leaded to an irregular spindle morphology (Fig. 4D-H, J and K, red arrows); 2), irregular assembly of microtubules surrounding chromosomes (Fig. 4D, H, L and M; Supplemental Fig. S1G, blue arrows); 3), aberrant alignment of chromosomes (Fig. 4N, yellow arrow). F-53B did not interfere with microtubule organization at anaphase I and/or interkinesis stages (Supplemental Fig. S1C in control; H in 50 μM F-53B, P and Q in 100 μM F-53B). At metaphase II, defective chromosome distribution was visualized in Col-0 exposed to 50 μM F-53B (Supplemental Fig. S1I in 50 μM and R in 100 μM F-53B, normal; Supplemental Fig. S1J, Fig. 4I, abnormal, yellow arrows). Under 100 μM F-53B exposure conditions, meiocytes at telophase II showing phragmoplasts with an altered orientation and sparse microtubules were observed (Fig. 4C in control, Supplemental Fig. S1K and L in 50 μM F-53B and S in 100 μM F-53B, normal; Fig. 4O, abnormal). At the end of meiosis, meiocytes under both growth conditions produced normal tetrads (Supplemental Fig. S1D, M and T). Overall, these data indicate that F-53B interferes with spindle and phragmoplast organization in Arabidopsis male meiosis.

### 3.5 F-53B reduces CO formation

The number and position of chiasmata, the chromosome structures that manifest the CO sites between homologs, can determine the configuration a bivalent (Huang et al., 2022). In general, the configurations of a bivalent include: a ‘O’-like shape determined by two chiasmata (Fig. 3A; i, green arrow); a ‘8’-like shape that indicates three chiasmata in a bivalent (Fig. 3A; ii, yellow arrow); a rod shape, in which homologs are linked by one distally-distributed chiasma (Fig. 3B and C; iii, red arrows); and a ‘X’-like shape (Fig. 3C; iv, blue arrow), which also indicates one chiasma in a bivalent (Huang et al., 2022). To confirm that F-53B influences CO formation, we first compared the configurations of bivalents in Col under control and F-53B exposure conditions. In control, 46.2% and 35.3% bivalents exhibited an ‘O’-like shape and an ‘X’-like shape, respectively, with the bivalents showing an ‘8’-like shape or a rod shape occurring at 7.7% and 10.8%, respectively (Fig. 4D). This composition pattern of bivalent configurations led to an average of nine chiasmata per meiocyte in Col-0 under control conditions (Fig. 4E). In Col-0 exposed to 50 μM F-53B, the bivalent showing an ‘8’-like configuration was not visualized (Fig. 4D), however, the fraction of the bivalent displaying a rod shape was largely increased (Fig. 4D, 41.7%). This alteration suggested that the proportion of bivalents with one chiasma was elevated. A similar composition of the configurations of bivalents was observed in Col-0 exposed to 100 μM F-53B. Specifically, the bivalents displaying a rod (27.5%) or an ‘X’-like (45.0%) shape occurred at higher levels than those in control (Fig. 4D). Accordingly, the numbers of chiasma in Col-0 exposed to 50 or 100 μM F-53B decreased to an average of 6.5 and 5.0 per meiocyte, respectively, both of which were significantly lower than the value in control (Fig. 4E; control vs. 50 μM F-53B, *P* < 0.001; control vs. 100 μM F-53B, *P* < 0.001). These data suggest that meiosis in Arabidopsis exposed to F-53B produces a reduced number of CO.

To consolidate that F-53B reduces CO formation, we performed immunolocalization of the type-I class CO recombinase HEI10 in PMCs and quantified its abundance on diakinesis chromosomes in Col-0 exposed to F-53B. In control PMCs, an average of 7.0 HEI10 foci per PMC was detected (Fig. 4F and I). In comparison, Col-0 exposed to 50 or 100 μM F-53B exhibited an average of 6.5 and 5.0 HEI10 foci, respectively, which were significantly less than that in control (Fig. 4G, H and I; for control vs. 50 μM F-53B, *P* < 0.05; for control vs. 100 μM F-53B, *P* < 0.001). Taken together, these data demonstrate that F-53B reduces type I CO formation in Arabidopsis.

### 3.6 Arabidopsis exposed to F-53B shows a reduced number of DMC1 protein foci on zygotene chromosomes

Because CO rate is positively correlated with the number of DSBs (Sidhu et al., 2015), and there was no obvious defect in homolog pairing observed in Col-0 exposed to F-53B, we hypothesized that F-53B may reduce CO rate by attenuating DSB formation. To this end, we performed immunolocalization of the key DSB repair protein DMC1, which catalyzes DSB repair using homologs as the template (Da Ines et al., 2013; Kurzbauer et al., 2012), on zygotene chromosomes and quantified its foci number. The meiosis-specific chromosome axis protein SYN1 (Bai et al., 1999; Cai et al., 2003) was co-immunostained, which was used to label the chromosomes and to indicate the developmental stage of the meiocytes. In control plants, we detected an average of 85.5 DMC1 foci per meiocyte (Fig. 6A and D). In comparison, Col-0 exposed to 50 or 100 μM F-53B showed an average of 55.0 and 43.5 DMC1 foci, respectively, which were significantly less than the value in control (Fig. 6B, C and D; for 50 μM F-53B, *P* < 0.05; for 100 μM F-53B, *P* < 0.01). These findings suggest that F-53B may either reduce the loading of DMC1 on zygotene chromosomes, or may reduce meiotic DSB formation, or both, which at least partially contributes to the lowered CO rate.

### 3.7 Arabidopsis *atm* mutant shows reduced meiotic chromosome fragments after the exposure to F-53B

To confirm that F-53B reduces DSB formation in Arabidopsis meiocytes, we applied a genetic approach by evaluating the sensitivity of meiotic chromosome integrity in Arabidopsis *atm* mutant to F-53B. The *atm* mutant has a defect in RAD51-meidated DSB repair and shows chromosome fragments during meiosis, the level of which is responsive to both endogenous genetic mutations and exogenous abiotic stresses that trigger an increased or reduced DSB formation (Kurzbauer et al., 2021; Zhao et al., 2023). Hence, by scoring the number of chromosome fragments in *atm* exposed to F-53B, it enables evaluation of the impact of F-53B on meiotic DNA stability. In our assay system, *atm* under control conditions yielded an average of 6.0 fragmented chromosome bodies per meiocyte at stages from metaphase I to telophase II (Fig. 5F-J, T). In comparison, *atm* exposed to 50 and 100 μM F-53B produced 4.5 and 3.0 fragments per meiocyte, respectively (Fig. 5K-S, T), the latter of which was significantly less than the value in *atm* under control conditions (Fig. 5T, P < 0.01). In contrast, Col under either control conditions or exposure to 50 or 100 μM F-53B did not produce any chromosome fragment (Fig. 5A-E, T). These data suggest that Arabidopsis exposed to F-53B produce a reduced number of meiotic DSBs.

**Figure 5.**
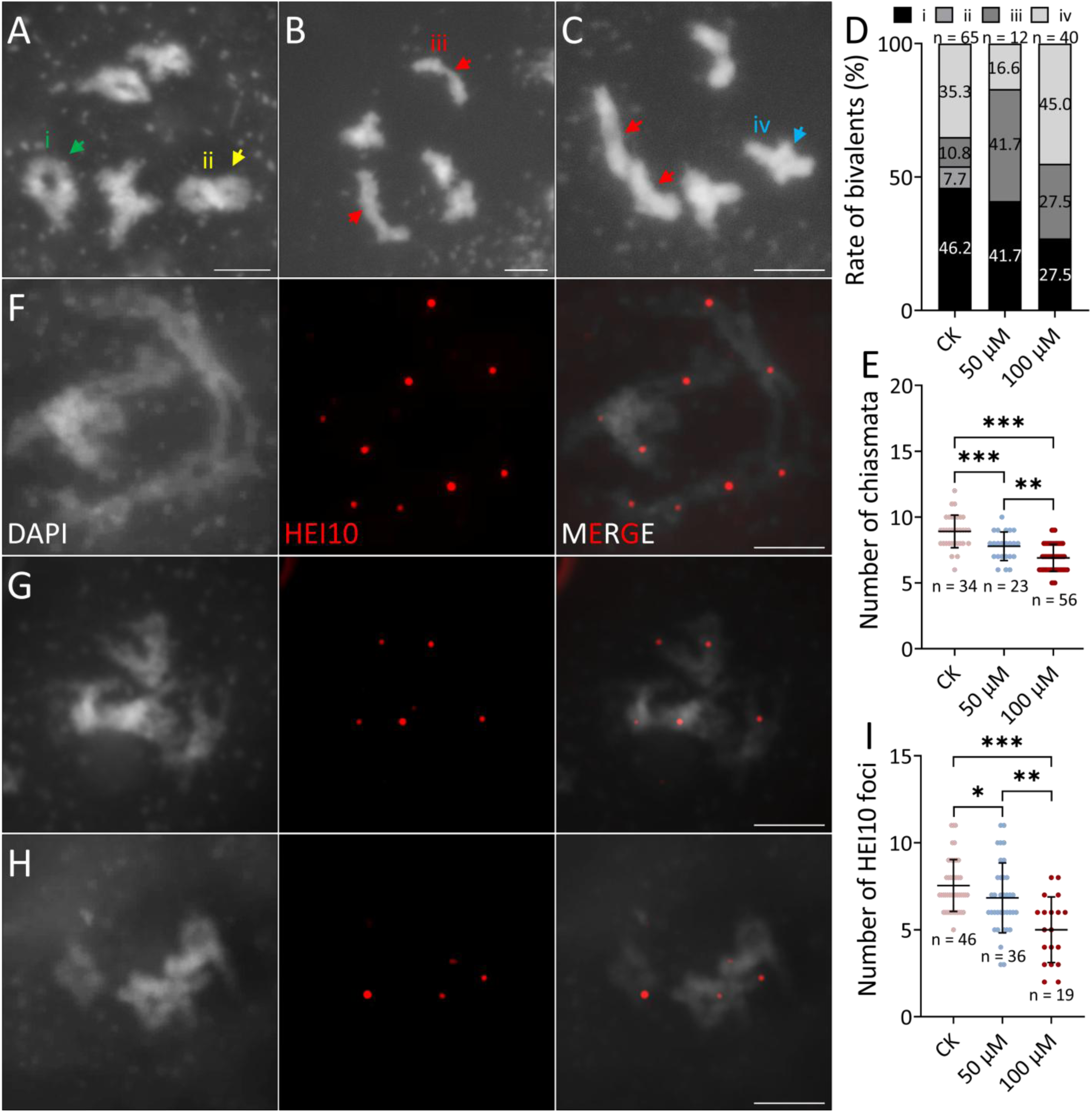
Evaluation of CO level in Arabidopsis exposed to F-53B. A-C, DAPI-stained meiocytes at diakinesis stage in Col-0 under control conditions (A) or exposed to 50 (B) or 100 μM (C) F-53B for 48 h. The green arrow indicates a bivalent showing an ‘O’-like shape; yellow arrow indicates a bivalent showing an ‘8’-like shape; red arrows indicate bivalents showing a rod shape; blue arrow indicates a bivalent showing an ‘X’-like shape. F-G, Immunolocalization of HEI10 on diakinesis chromosomes in Col-0 under control conditions (F) and exposed to 50 (G) or 100 μM (H) F-53B for 48 h. Scale bars = 5 μm. D, E and I, Graphs showing the composition of bivalent configurations (D), the number of chiasmata on diakinesis chromosomes (E) and the number of HEI10 foci on diakinesis chromosomes (I) in Col-0 under control conditions, or exposed to 50 or 100 μM F-53B for 48 h. Significance levels were determined based on unpaired *t* tests; *** indicates *P* < 0.001; ** indicates *P* < 0.01; * indicates *P* < 0.05; in panel D, the rates of the bivalents showing the corresponding configurations are shown; n indicates the number of analyzed bivalents (D) or meiocytes (E and I).

**Figure 6.**
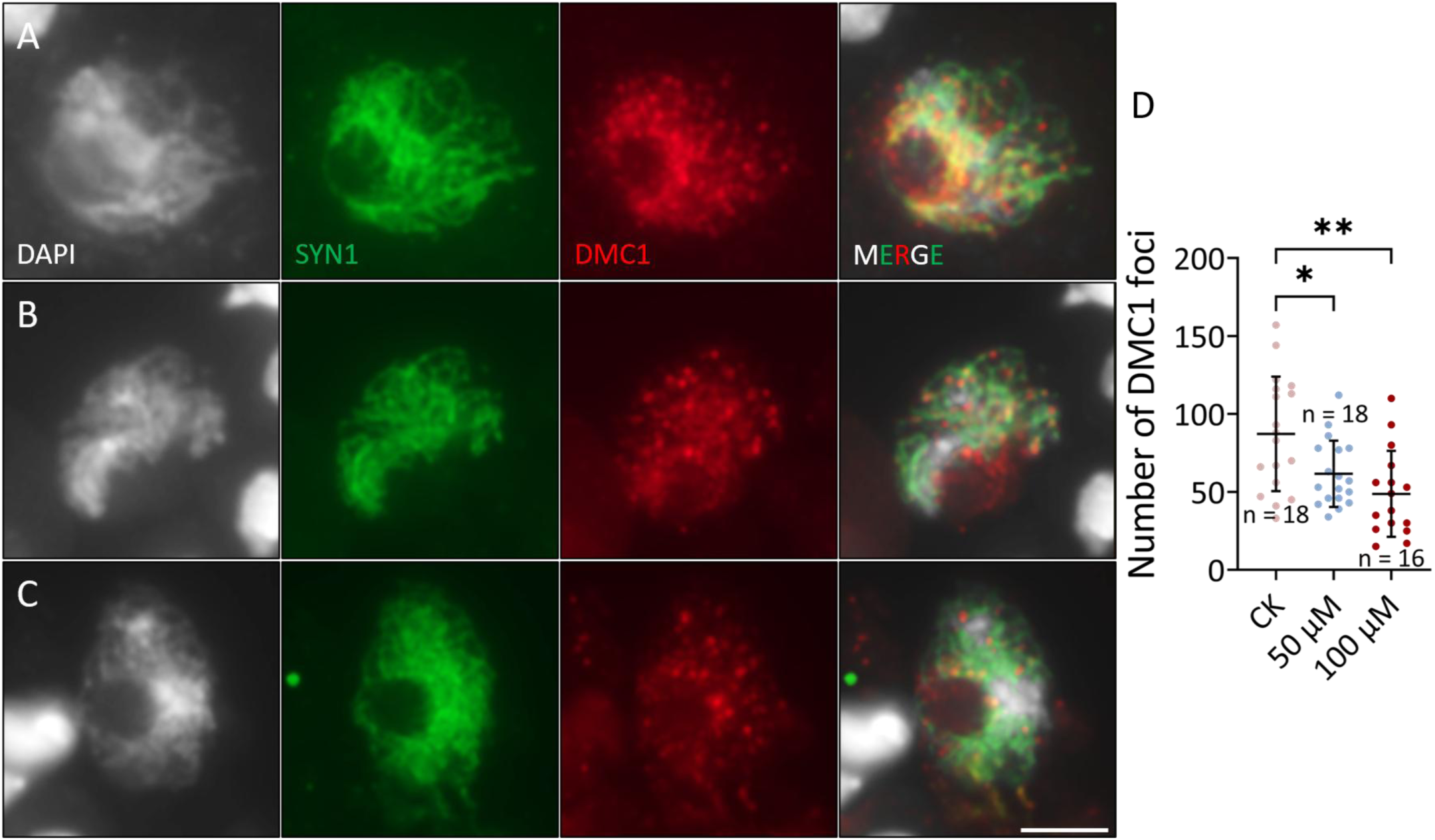
F-53B reduces the number of DMC1 foci on zygotene chromosomes. A-C, Co-immunolocalization of SYN1 and DMC1 on zygotene chromosomes in Col-0 under control conditions (A), or exposed to 50 (B) or 100 μM (C) F-53B. D, Graph showing the number of DMC1 foci on zygotene chromosomes in Col-0 under control conditions or exposed to 50 or 100 μM F-53B. Significance levels were determined based on unpaired *t* tests; ** indicates *P* < 0.01; * indicates *P* < 0.05; n indicates the number of analyzed meiocytes; the scale bar applies to all panels in this figure and indicates 5 μm.

**Figure 7.**
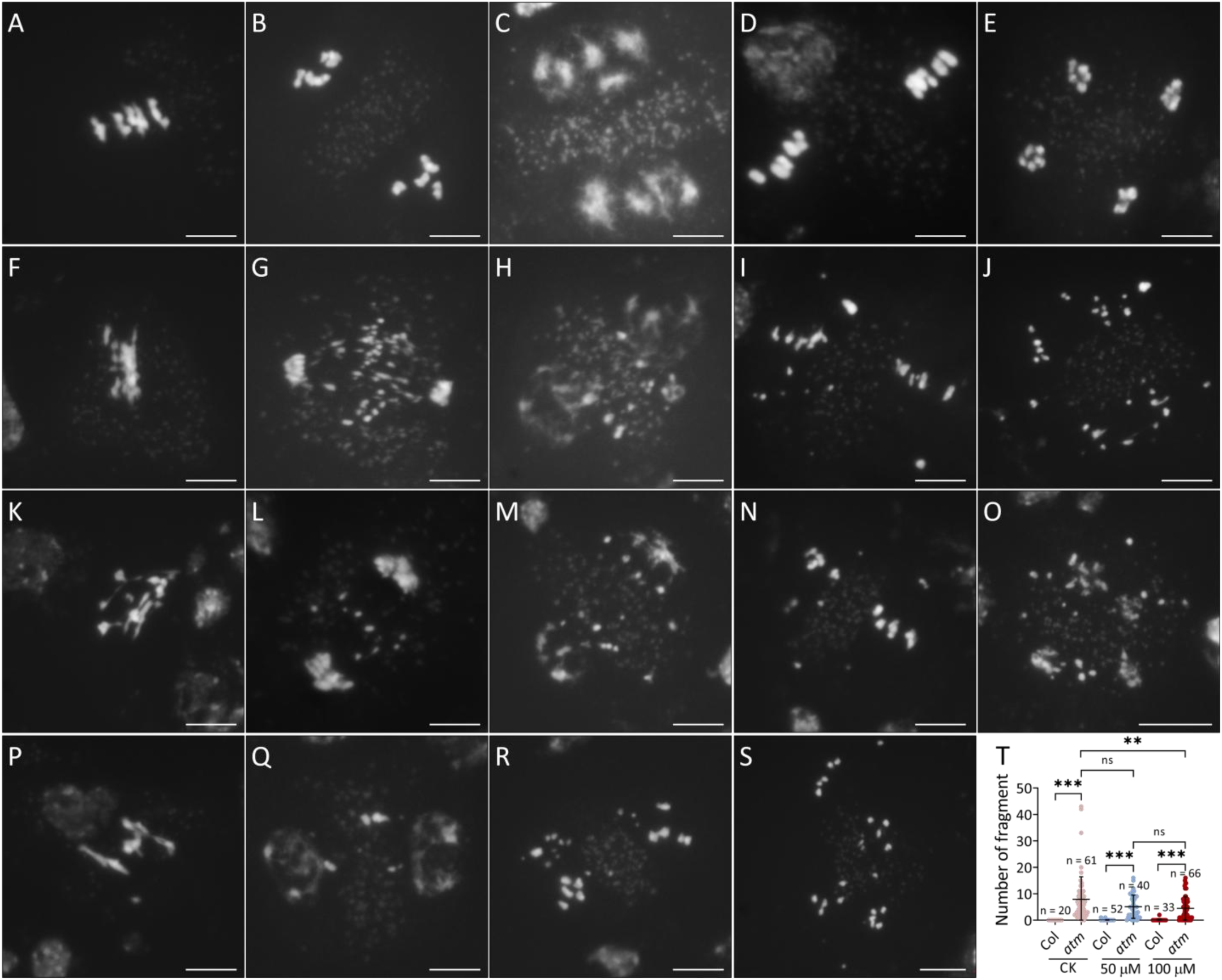
F-53B reduces meiotic chromosome fragments in *atm*. A-E, Representative DAPI-stained chromosome spreads meiocytes at metaphase I (A), anaphase I (B), interkinesis (C), metaphase II (D) and telophase II (E) stages in Col-0 under control conditions, or exposed to 50 or 100 μM F-53B. F-S, DAPI-stained chromosome spreads in *atm* meiocytes at metaphase I (F, K and P), anaphase I (G and L), interkinesis (H, M and Q), metaphase II (I, N and R) and telophase II (J, O and S) stages under control conditions (F-J), or exposed to 50 (K-O) or 100 μM (P-S) F-53B. Scale bars = 5 μm. T, Graph showing the number of chromosome fragments in Col-0 (‘Col’ is used here for simplification) and *atm* under control conditions, or exposed to 50 or 100 μM F-53B. Significance levels were determined based on unpaired *t* tests; *** indicates *P* < 0.001; ** indicates *P* < 0.01; ns indicates *P* > 0.01; n indicates the number of analyzed meiocytes.

## 4 Discussion

In this study, we showed that F-53B interferes with meiotic cell division and disrupts anther and embryo development in Arabidopsis, which reveals that as the cases in mammals (Feng et al., 2024; He et al., 2022; Qiu et al., 2024; Wang et al., 2024; Wu et al., 2023; Zhang et al., 2025), F-53B is toxic to both somatic and reproductive cells in plants (Ebinezer et al., 2022; Li et al., 2023; Pan et al., 2021; Zhao et al., 2025). Arabidopsis exposed to F-53B showed an aborted or disrupted embryo development resulting in impaired seed setting. The undeveloped ovules with small sizes indicate defects in fertilization, which can be attributed to an impaired gametogenesis and the production of viable male and female gametes. This notion can be supported by the observation of a disruption of anthers in F-53B-exposed plants. The impaired expression of AMS in the anthers could be owing to F-53B-induced unviability of the tapetum which has been evidence to be hypersensitive to abiotic stresses (Parish et al., 2012; Zhao et al., 2025). During gametogenesis tapetal cells undergo PCD, the occurrence and timing of which are controlled by a dynamic alteration of the ROS level in the anthers (Yi et al., 2016; Yu et al., 2017; Zheng et al., 2019). Despite a divergence of the impact of F-53B on ROS induction between species (Amstutz et al., 2025; Li et al., 2023; Lin et al., 2020; Pan et al., 2021; Yang et al., 2023; Zhao et al., 2025), it is still possible that F-53B evokes an ectopic alteration in the ROS level dynamics in the anthers which triggers premature occurrence of PCD and degradation of the tapetum.

We showed in this study that F-53B interferes with chromosome distribution and/or arrangement in meiosis I and II. Because no obvious defects were found in balanced segregation of homologous chromosomes and sister chromatids, we propose that the function of the microtubule cytoskeleton in Arabidopsis meiocytes is not suppressed but is only disturbed by F-53B. In support, the assembly of the spindle and phragmoplast structures can occur in F-53B-exposed meiocytes, but their configuration and orientation are interfered, which could lead to the lesions in arrangement and alignment of the chromosomes. Our findings reveals a conservation of the impact of F-53B on meiotic chromosomes and microtubule organization in mouse oocytes, in which F-53B was shown to induce defective spindle organization and chromosome alignment (Chu et al., 2025). The configuration of multi-axial spindles in F-53B-exposed meiocytes mimicked the phenotypes observed in Arabidopsis with dysfunction of microtubule-based motor protein ATK1, which typically hydrolyzes ATP to generate a motive force along a microtubule (Chen et al., 2002; Hotta et al., 2022). Chu et al. reported that F-53B induces abortion of meiosis progression and the spindle defects by disrupting mitochondria cellular distribution (Chu et al., 2025). We thus speculate that in both mammals and plants, F-53B may damages the energy-supply system that is crucial for the localization and/or function of microtubule polymerization regulators in meiocytes (Chu et al., 2025; Gao et al., 2017; Wan et al., 2020). Considering that F-53B has been shown to affect the organization of microtubules and actin filament and induce aberrant cell morphology in somatic cells (Gao et al., 2017; Wan et al., 2020; Zhao et al., 2025), it is likely that cytoskeleton network is a prominent target of F-53B.

In both mammals and plants, it was unclear whether F-53B has an impact on meiotic recombination that is the basis for biological diversity. In this study, we showed that F-53B reduces chiasmata and HEI10 foci formation on diakinesis chromosomes, which suggests that F-53B attenuates CO rate in plants. The reduction levels of chiasmata and the HEI10 foci occurred at a similar level, implying that F-53B primarily lowers type I CO formation. The occasionally visualized univalents suggest that the function and/or stability of the regulators involved in CO assurance are interfered by F-53B. Since the configuration of chromosomes at pachytene appeared normal, F-53B-induced reduction of CO does not likely result from defects in homolog pairing, but is possibly owing to an interfered expression and/or function of the ZMM recombinases, e.g. MLH1 and HEI10. Alternatively, it is also possible that the compromised CO formation is partially caused by a reduced DSB abundance, which is positively correlated with CO level (Xue et al., 2018). In support of this hypothesis, we showed that F-53B reduces the number of DSB repair protein DMC1 on zygotene chromosomes and it promotes the meiotic chromosome integrity in *atm*. These findings together with our previous report reveal that the toxic effect of F-53B on DNA stability varies between species or cell types (Li et al., 2023; Pan et al., 2021; Qiu et al., 2024; Zhao et al., 2025). Taken together, our study reveals an impact of F-53B on meiotic recombination and thus on genetic makeup of gametes. At a broader perspective, considering the global distribution and accumulation of F-53B in various environment matrix, the toxicity of F-53B to sexual cells in plants highlights its potential threat to ecological diversity and evolution.

Furthermore, some limitations exist in this study. We assessed the toxicity of F-53B on male reproductive development in Arabidopsis plants cut with main shoots, an exposure method based on the transpiration flow in the inflorescences (Armstrong, 2013). Although this method has been widely used in determining the time-course and other features of meiosis, based on which some theoretical models have been built (Hernández Sánchez-Rebato et al., 2024; Sanchez-Moran et al., 2007), the exposure system applied here may not reflect a practical environmental damage or level that F-53B may cause. Nevertheless, despite these limitations, our study provides cytogenetic insights into toxicity of F-53B to the reproductive cells in plants. Studies in crops, which are naturally exposed to contaminated lands, are needed in future to evaluate and decipher the accumulation and the toxicity of F-53B in plant reproductive system.

## CRediT authorship contribution statement

Y.C., X.C., Z.R., H.F., Y.L., L.X. and Z.S. performed the investigation; Y.Q., G.Y. and X.L. contributed to data analysis; B.L. conceived project, analyzed data, wrote and edited the manuscript. All authors have agreed with the manuscript.

## Declaration of Competing Interest

The authors declare no conflicts of interest.

## Data Availability

The data that support the findings of this study are available from the senior author (B.L.) upon reasonable request.

## Acknowledgements

This study was supported by National Natural Science Foundation of China (22206127 to X.L.; 32572136 to Z.R; 32270364 and 31270361 to Y.G.), the Program of Science and Technology Commission of Shanghai Municipality (21ZR1435800 to X.L.), the Fund for Scientific Research Platforms of South-Central Minzu University (PTZ25018 to Y.Q.), Hubei Provincial Natural Science Foundation (2024AFB695 To B.L.) and the Fundamental Research Funds for the Central Universities, South-Central Minzu University (CZZ24011 to B.L.).

## Appendix A. Supporting information

Supplementary data associated with this article can be found in the online version

## Notes

### Competing Interest Statement

The authors have declared no competing interest.

