## Supplemental file for "The emerging contaminant Cl-PFESA/F-53B is toxic to meiotic cell division and reproduction in *Arabidopsis thaliana*"

Running title: F-53B interferes with meiosis in plants

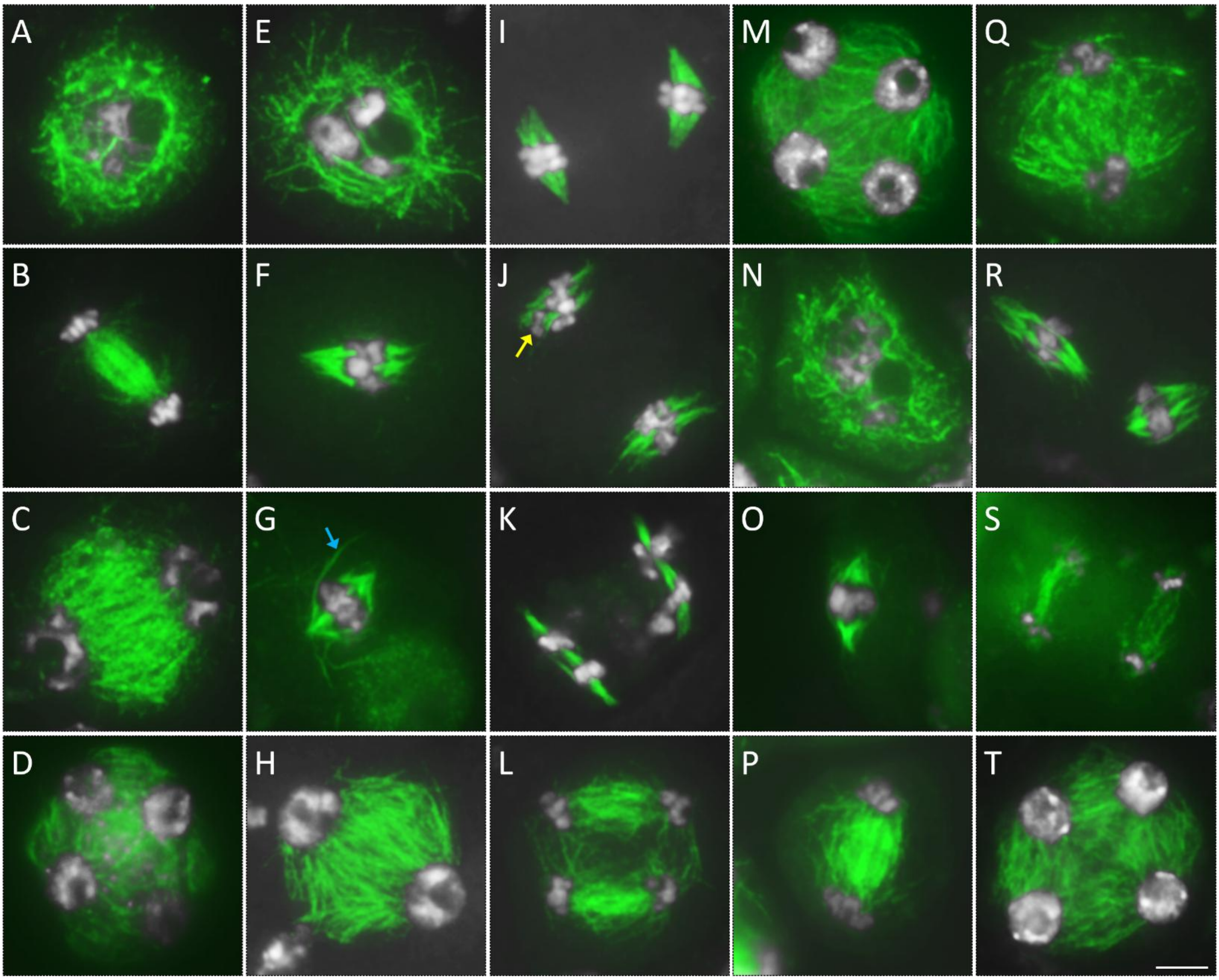


Supplemental Figure S1. Arabidopsis exposed to F-53B shows alterations in microtubule organization during male meiosis. A-T, Immunolocalization of α-tubulin in meiocytes at diakinesis (A, E and N), metaphase I (F, G and O), anaphase I (B and P), interkinesis (C, H and Q), metaphase II (I, J and R), telophase II (K, L and S) and tetrad (D, M and T) stages in Arabidopsis under control conditions (A-D), or exposed to 50 (E-M) or 100 μM (N-T) F-53B. White, DAPI; green, α-tubulin. The blue arrows indicate irregularly-assembled microtubule filaments; yellow arrows indicate defectively-distributed chromosomes. The scale bar applies to all panels in this figure and indicates 5 μm.

Supplemental Table S1. Primers used in this study.

| Primers | Sequence (5’ - 3’) | Purpose |
|---|---|---|
| atm-2 F | ATCCATGTGGTTCAGTCTTGC | *atm-2* genotyping |
| atm-2 R | TTGGTATCCTGCAGAGGAAAG | |
